# Selective life-long suppression of an odor processing channel in response to critical period experience

**DOI:** 10.1101/2025.07.18.665601

**Authors:** Hans C. Leier, Julius Jonaitis, Alexander J. Foden, Abigail J. Wilkov, Paola Van der Linden Costello, Heather T. Broihier, Andrew M. Dacks

## Abstract

Sensory circuits undergo experience-dependent plasticity during early-life critical periods, attuning the nervous system to levels of key environmental stimuli. During a critical period in the *Drosophila* olfactory system, we found that exposure to ethyl butyrate (EB) induces glial phagocytosis of odorant receptor Or42a-positive olfactory sensory neuron (OSN) axon terminals which terminate in the VM7 glomerulus (Leier and Foden et al., 2025). Here, we extend these findings by establishing functional significance and circuit selectivity in this critical period paradigm. First, using a combination of two-photon Ca^2+^ imaging and the genetically-encoded voltage indicator ASAP5, we find that Or42a OSN odor-evoked responses are permanently suppressed in animals with critical period odor exposure. Thus, critical period odor exposure results in long-term changes to odor sensitivity in Or42a OSNs. Second, to establish the selectivity of glial pruning for Or42a axon terminals, we examined projection neurons (PNs) postsynaptic to Or42a OSNs as well as a second population of highly EB-responsive OSNs, called Or43b OSNs. We find that (1) within VM7, glial pruning is selective for Or42a terminals, and (2) while Or43b OSNs appear modestly pruned, they maintain their sensitivity to EB. To elucidate this difference, we turned to the *Drosophila* connectome. We identify striking differences in the scale of inhibitory connectivity to Or42a and Or43b OSNs, suggesting that Or42a OSNs may play a particularly central role in EB odor processing. This study expands our understanding of this critical period plasticity paradigm by demonstrating life-long suppression of pruned Or42a OSNs and establishing its specificity within and between sensory circuits.

## Introduction

Animals integrate information from multiple sensory processing channels to seek out food, reproduce, and avoid danger. In the olfactory system, sensory channels are defined by populations of first-order olfactory sensory neurons (OSNs) expressing unique repertoires of chemoreceptor proteins. For many species, these parallel odor-processing channels converge at an initial processing stage (the olfactory bulb in mammals and the antennal lobe (AL) in insects) in which all OSNs expressing the same combination of receptors synapse at stereotyped glomeruli onto local interneurons and second-order olfactory neurons that project to other regions of the brain (Schlegel et al., 2021; Wilson, 2013).

Because many odorant receptors (ORs) are tuned to recognize a range of ligands with similar structures (Grabe et al., 2016), and many ethologically relevant odorants can bind multiple classes of ORs with varying affinity (Hallem et al., 2004; Hallem and Carlson, 2006; Knaden et al., 2012; Schubert et al., 2014), olfactory circuits are capable of combinatorially encoding a vast odor space of unique compounds with a comparatively small number of receptors (Si et al., 2019). This remarkable capacity is observed even in relatively simple olfactory systems, such as *Drosophila melanogaster*’s repertoire of ∼60 OR genes expressed across ∼2,600 OSNs (Benton et al., 2025).

While this method of odor coding far exceeds the discriminatory capacity of other sensory systems (Bushdid et al., 2014), it also presents a major vulnerability: at sufficiently high levels, a so-called ‘public odor’ capable of activating many classes of OSNs could exceed the circuit’s ability to discriminate between similar compounds, rendering large parts of the odor landscape inaccessible (Hallem and Carlson, 2006). To counteract this, the *Drosophila* AL features multiple adaptations. For instance, local interneurons (LNs) synapse with both OSNs and second-order projection neurons (PNs) and exert gain control over the OSN-to-PN relay, using both lateral excitation and inhibition to maintain odor coding across orders of magnitude of intensity (Barth-Maron et al., 2023; Liu and Wilson, 2013; Olsen et al., 2010; Olsen and Wilson, 2008; Root et al., 2008, 2007). Gain control allows recent network activity to impact odor-evoked responses over short time periods, whereas prolonged odorant exposure can induce larger scale circuit remodeling, a phenomenon largely restricted to early-life windows of heightened plasticity (Chodankar et al., 2020; Das et al., 2011; Devaud et al., 2003, 2001; Golovin et al., 2019; Gugel et al., 2023; Jindal et al., 2023; Mallick et al., 2024; Sachse et al., 2007; Sudhakaran et al., 2012).

Recently, we and others uncovered the cellular mechanism of ethyl butyrate (EB)-induced circuit remodeling in the glomerulus VM7, which is innervated by OSNs expressing the EB-sensitive OR Or42a (Leier et al., 2025; Nelson et al., 2024). EB is an important food cue detected by large numbers of ORs, with Or42a being among the most sensitive (Hallem et al., 2004; Hallem and Carlson, 2006; Semmelhack and Wang, 2009). During a 48-hour critical period following eclosion, we found that EB exposure induces glia to invade VM7 and phagocytose Or42a OSN axon terminals, drastically reducing OSN-PN connectivity even after several days had passed (Leier and Foden et al., 2025). From a circuit standpoint, we hypothesized that this may be an alternative method of limiting input from the most active information channel in an over-stimulating environment, preserving odor-coding capacity of other EB-sensitive OSNs (Tadres et al., 2022).

In the current study, we define functional consequences and cellular specificity in this critical period paradigm. We previously found that early odor exposure led to a reduction of spontaneous activity of VM7 PNs (Leier and Foden, 2025), but we did not probe changes in odor-evoked activity. Here, employing two-photon Ca^2+^ imaging and a genetically-encoded voltage indicator, we find that following critical period odor exposure, Or42a OSN odor-evoked responses remain suppressed at 25 days post eclosion (DPE). These data indicate that critical period EB exposure permanently alters odor processing.

We next addressed if glial pruning is restricted to Or42a OSN axon terminals or whether other neurons engaged in EB odor processing are likewise pruned. Strikingly, within the Or42a-VM7 PN circuit, pruning is selective for Or42a axon terminals since neither axons nor dendrites of VM7 PNs are refined following critical period odor exposure. We then turned to VM2, the second-most EB sensitive glomerulus in *Drosophila*, which is innervated by OSNs expressing Or43b (Olsen et al., 2007). While Or43b OSNs are modestly pruned following early-life EB exposure, their odor-evoked responses are unchanged. The existence of critical period Or43b remodeling supports our working hypothesis that pruning of exquisitely EB-sensitive OSNs dampens activity in these hyperactive information channels to maintain odor coding capability. However, we were surprised that EB-evoked responses of Or43b neurons remain relatively normal, arguing that pruning of Or43b presynaptic terminals does not attenuate odor responses below baseline. To elucidate distinct functional consequences of Or42a and Or43b pruning, we capitalized on the recently completed *Drosophila* connectome (Dorkenwald et al., 2024; Schlegel et al., 2024; Zheng et al., 2018) and compared connectivity of Or42a and Or43b neurons. Our connectomic analysis revealed large-scale differences in the local inhibitory circuitry interacting with OSNs within VM7 and VM2. Specifically, while there are roughly comparable numbers of Or42a and Or43b OSNs, Or42a neurons make roughly seven-fold more synapses with a specific class of highly connected inhibitory interneurons, suggesting Or42a neurons serve an outsize function in EB odor processing. Together, we propose that glial pruning and synaptic inhibition play complementary roles in an interlocking mechanism to adapt and maintain the odor-coding capacity of the olfactory system.

## Results and Discussion

### Life-long suppression of Or42a OSNs after critical-period EB exposure

We previously showed using whole-cell patch electrophysiological recordings from PNs (Figure 1A) that spontaneous release from Or42a OSN axon terminals is reduced by ∼98% following critical-period exposure to EB and remains similarly depressed after a 4-day recovery period (Leier and Foden et al., 2025). While this *ex vivo* recording strategy validated the loss of presynapses observed by confocal microscopy, it did not address the effects of critical-period exposure on the odor-evoked responses of Or42a OSNs. To address this, we used two-photon microscopy to measure Ca^2+^ flux in Or42a OSNs expressing GCaMP8f (Zhang et al., 2023), following 0–2 DPE 15% EB exposure (Figure 1B).

**Figure 1.**
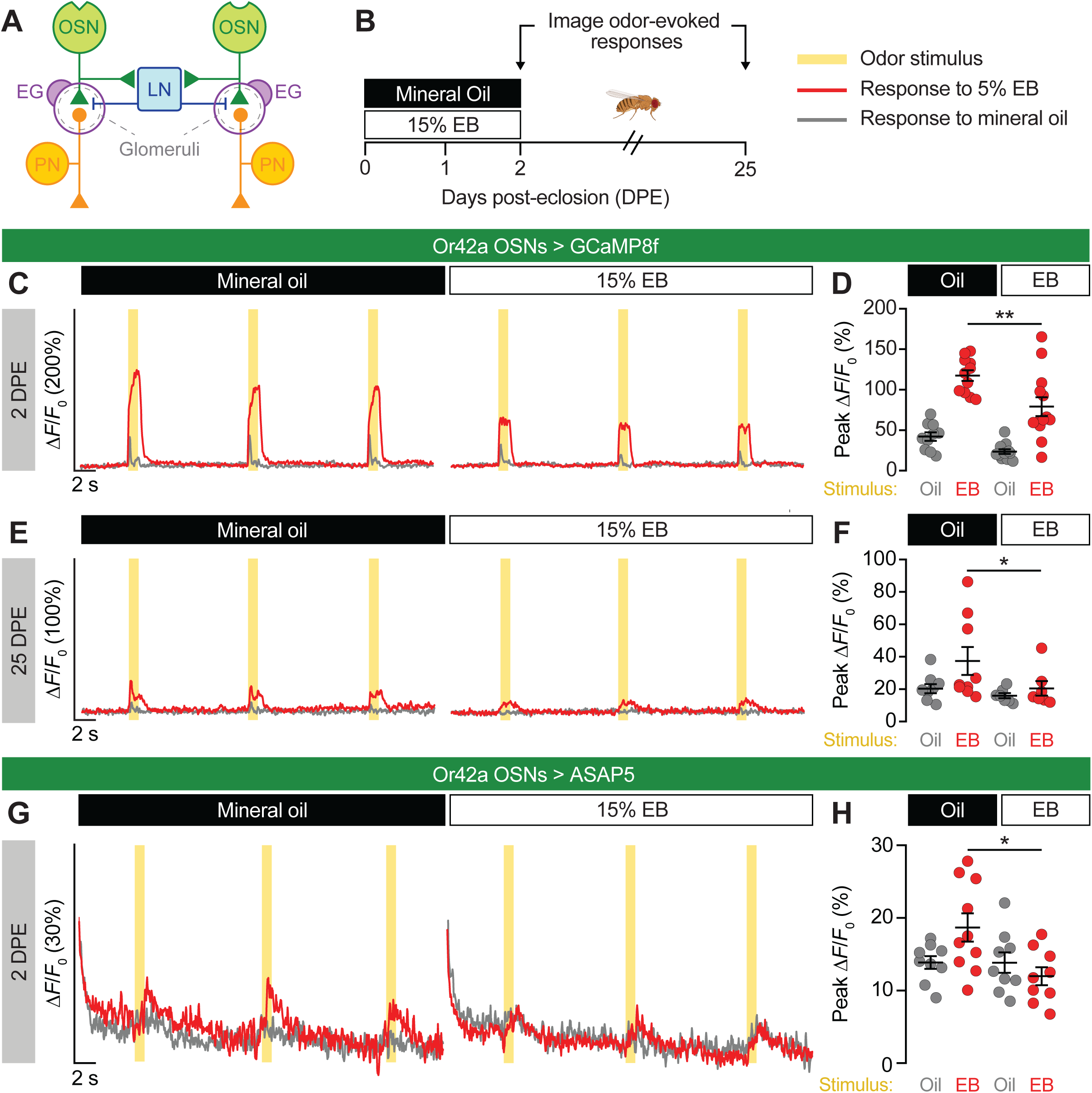
Lifelong suppression of Or42a OSNs after critical-period EB exposure. (**A**) Schematic of the *Drosophila* olfactory circuit at the level of the AL. OSN, olfactory sensory neuron, PN, projection neuron, LN, local interneuron, EG, ensheathing glia. (**B**) Overview of odorant-exposure experiments. Flies are exposed to 15% (v/v) ethyl butyrate (EB) or mineral oil vehicle control during the Or42a OSN critical period from 0–2 days post-eclosion (DPE). (**C**–**F**) Live imaging of Or42a OSN odor-evoked activity at 2 (**C**, **D**) or 25 (**E**, **F**) DPE, following exposure to 15% EB or mineral oil from 0–2 DPE. Or42a-GAL4>UAS-GCaMP8f flies were stimulated with three 1-second puffs (yellow bars) of 5% EB (red traces and data points) or mineral oil vehicle control (grey traces and data points) spaced 10 seconds apart. Traces (**C**, **E**) show the mean ΔF/F_0_ values of all trials. Data points (**D**, **F**) represent the mean peak ΔF/F_0_ values for each fly. (**G**–**H**) Live imaging of ASAP5 responses in Or42a OSNs in 2 DPE flies following exposure to 15% EB or mineral oil from 0–2 DPE. Flies were stimulated with odor puffs as above. Traces (**G**) represent the mean ΔF/F_0_ values of all trials. Data points (**H**) represent the mean peak ΔF/F_0_ values for each animal. Data in **D**, **F**, and **H** are mean ± SD. *p < 0.05, **p < 0.01, unpaired t test (**D**) or Mann-Whitney U-test (**F**, **H**). Genotypes, raw values, and detailed statistics are provided in Figure 1—source data 1.

In VM7, odor-evoked OSN responses were sharply reduced in flies that had been exposed to EB, relative to mineral oil-exposed controls (Figure 1C–D). Curious about how long this effect persisted, we aged 0–2 DPE-exposed flies to 25 DPE and repeated our imaging. Remarkably, even at an advanced age where olfactory function has declined considerably (Hussain et al., 2018), odor-evoked activity remained suppressed in flies that had been exposed to EB nearly a month earlier (Figure 1E–F). To our knowledge this is the longest span over which early-life *Drosophila* olfactory circuit remodeling has been assessed, and highlights the remarkably permanent effects of glial pruning.

While live calcium imaging using GCaMP has been extensively used to study OSN activity in the *Drosophila* AL (Das et al., 2011; Hong and Wilson, 2015; Semmelhack and Wang, 2009; Sudhakaran et al., 2012), we considered whether the well-known limitations of the technique (Yang and St-Pierre, 2016) may be exacerbated when attempting to assess circuit activity in axon terminals that are actively being phagocytosed. To address this concern, we turned to the recently developed genetically encoded voltage indicator (GEVI) ASAP5 (Hao et al., 2024), using it for the first time in the *Drosophila* olfactory system. As with GCaMP, Or42a OSNs expressing ASAP5 displayed reduced odor-evoked activity after 0–2 DPE EB exposure compared to mineral oil controls (Figure 1G–H). While odor-evoked GEVI transients likely reflect both Ca^2+^ influx and action potentials at the axon terminal, the reduction in odor evoked activity (from either indicator) combined with the reduction in pre-synaptic markers in OSN terminals and the reduction in spontaneous EPSPs recorded from PNs (Leier and Foden et al., 2025) are all consistent with EB exposure dramatically reducing the input from Or42a OSNs to the AL.

### Experience-dependent pruning in VM7 is restricted to the presynaptic compartment

Thus far, our work has focused on the presynaptic side of the OSN-PN relay. There are numerous reports of early-life EB exposure inducing structural remodeling of PN dendrites in multiple glomeruli (Chodankar et al., 2020; Das et al., 2011; Golovin et al., 2019; McCann et al., 2011; Sachse et al., 2007), prompting us to carefully examine whether structural or physiological changes in PNs occur within VM7 during its critical period. We began by measuring the odor-evoked responses of VM7 PN dendrites, taking advantage of a newly characterized split-GAL4 line (Xie et al., 2021) to drive expression of GCaMP8f in just two cell types: VM7 and VM5 PNs. VM7 PN EB-evoked responses were reduced after critical-period EB exposure (Figure 2A–B), with a mean change at 2 DPE similar in magnitude to that of Or42a OSNs (45.4% for PNs vs. 32.9% for OSNs in Figure 1D).

**Figure 2.**
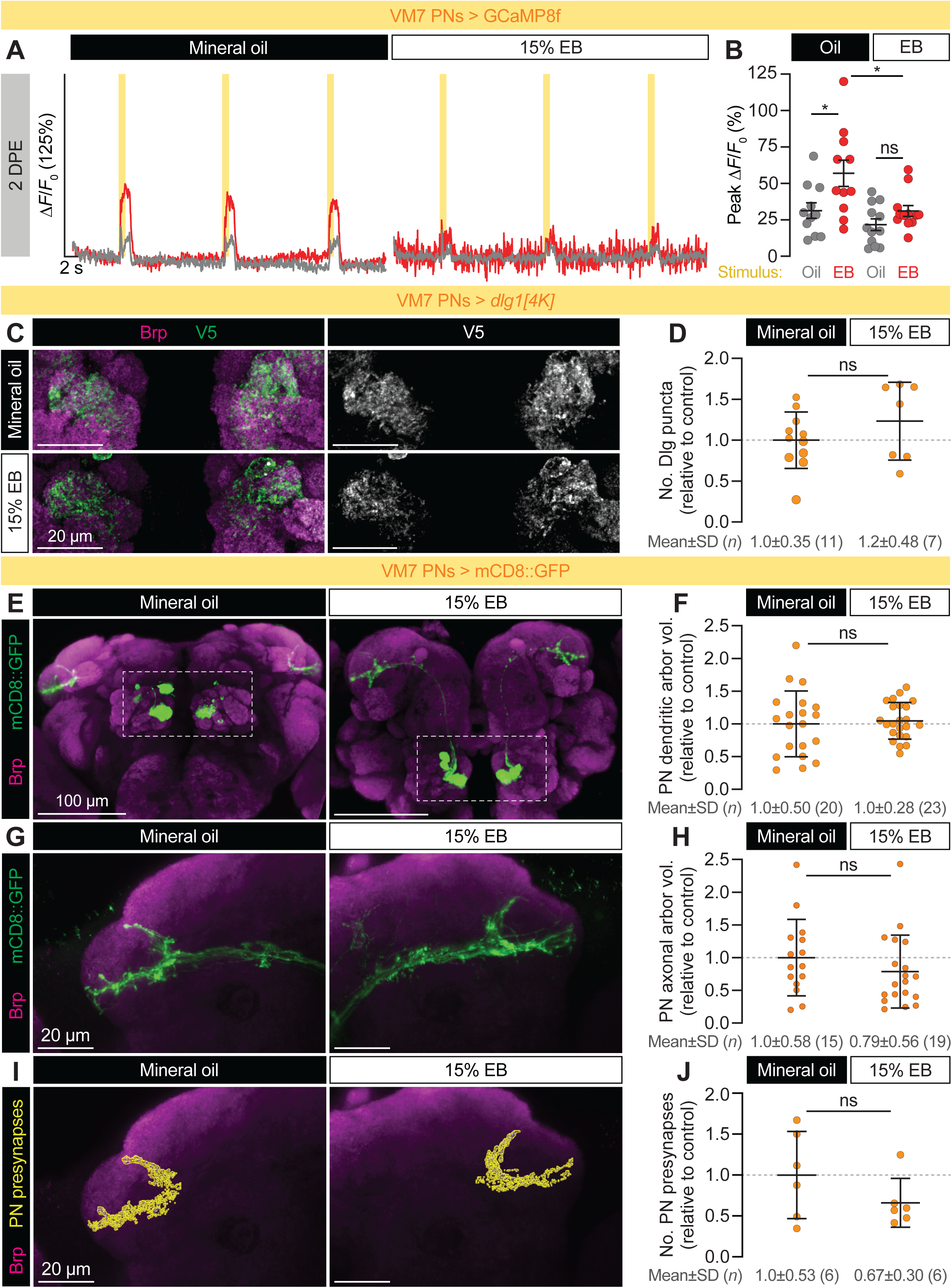
Experience-dependent pruning in VM7 is restricted to the presynaptic compartment. (**A**–**B**) Live imaging of VM7 PN odor-evoked activity at 2 DPE, following exposure to 15% EB or mineral oil from 0–2 DPE. VM7 PN split-GAL4>UAS-GCaMP8f flies were stimulated with three 1-second puffs (yellow bars) of 5% EB (red traces and data points) or mineral oil vehicle control (grey traces and data points) spaced 10 seconds apart. Traces (**A**) show the mean ΔF/F_0_ values of all trials. Data points (**B**) represent the mean peak ΔF/F_0_ values for each fly. (**C**) Representative dendritic arbors of VM7 PNs endogenously expressing the V5-tagged postsynaptic marker Discs large 1 (Dlg1), from flies exposed to mineral oil or 15% EB from 0–2 DPE. (**D**) Normalized counts of Dlg1 puncta. (**E**) Representative images of VM7 PNs after exposure to mineral oil or 15% EB from 0–2 DPE. Neuropil is visualized with antibody staining for the presynaptic active zone protein bruchpilot (Brp). PN dendritic arbors occupying VM7 are enclosed in white boxes. (**F**) Volumetric measurements of VM7 PN dendritic arbors shown in **E**. (**G**) Representative images of VM7 PN axonal arbors after exposure to mineral oil or 15% EB from 0–2 DPE. (**H**) Volumetric measurements of VM7 PN axonal arbors shown in **G**. (**I**) Models of VM7 PN presynapses from **G** after exposure to mineral oil or 15% EB from 0–2 DPE, defined as Brp puncta masked by mCD8::GFP-labeled VM7 PNs. (**J**) Normalized counts of VM7 PN presynapses from **I**. Data are mean ± SD. ns, p ≥ 0.05, *p < 0.05, Mann-Whitney U- test. Genotypes, raw values, and detailed statistics are provided in Figure 2—source data 1.

Was this loss in PN responses simply due to reduced innervation from their pruned presynaptic inputs, or had they been modified in other ways? Postsynaptic modifications play an important role in experience-dependent plasticity (Ehrlich and Malinow, 2004), including in the *Drosophila* olfactory system (Pribbenow et al., 2022); we therefore sought to visualize postsynaptic structures in VM7. To accomplish this, we used a newly developed tool to label excitatory postsynapses in a cell type-specific manner, wherein the endogenous postsynaptic scaffolding protein *discs large* (Dlg1; homolog of mammalian PSD-95) is tagged with V5 in cells expressing GAL4 (Parisi et al., 2023; Sheng and Kim, 2011). Using the same PN split-GAL4 driver as before, we expressed Dlg1-V5 in VM7 PNs and measured the number of postsynaptic puncta within VM7 (Figure 2C), finding that 15% EB exposure from 0–2 DPE did not alter VM7 PN postsynaptic content (Figure 2D). While this result may appear incongruous to the loss in presynapses seen in Or42a OSNs, compensation from other glomeruli may preserve the number of postsynapses in PNs (Olsen et al., 2007).

Also unlike their presynaptic partners, VM7 PN dendrites did not change in volume after 15% EB exposure from 0–2 DPE compared to mineral oil-exposed controls (Figures 2E, F) and they showed a striking absence of the punctate membrane appearance associated with glial phagocytosis of presynaptic axon terminals (Leier and Foden et al., 2025). Taken together, these data reinforce our previous finding that the intrinsic biophysical properties of VM7 PNs are unaltered even as they receive less innervation from pruned OSNs, and suggest that experience-dependent pruning is restricted to the presynaptic side of the VM7 OSN-PN relay.

Next, we wondered if VM7 PN axon terminals might be pruned or remodeled where they terminate in the mushroom body (Schlegel et al., 2021). While experience-dependent plasticity at PN axon terminals has been observed in other contexts (Kremer et al., 2010), less is known about their dynamics compared to plasticity of PN dendrites. With our VM7/VM5 PN split-GAL4 driving CD8::GFP, we were able to trace 4-5 PN axons from each AL to a tightly defined axonal arbor in the mushroom bodies (Figure 2G). As with PN dendrites, we found no volumetric changes in the axonal arbors of EB-exposed PNs (Figure 2H) and no changes in their presynaptic content (Figure 2I–J).

### Experience-dependent pruning and inhibition are decoupled in Or43b OSNs

Another outstanding question from our study of VM7 was whether early-life exposure has a uniform effect on the odor-evoked responses of OSNs in all glomeruli that respond to a single odor. EB robustly activates several classes of ORs in addition to Or42a (Figure 3A) (Münch and Galizia, 2016), making it well-suited to interrogate the extent of pruning in glomeruli throughout the AL. Of particular interest to us was VM2, which is innervated by OSNs expressing the odorant receptor (Or43b) with the second-highest affinity for EB (Münch and Galizia, 2016); indeed, Or43b and Or42a as a pair have significantly higher EB responses than other ORs (Figure 3A). Despite this, 0–2 DPE 15% EB exposure only caused a 12% reduction in Or43b OSN presynapses (Figure 3B) and an 18% reduction in VM2 volume (Figure 3C), far less than the changes seen in VM7. This relatively modest result suggests that, as a potential mechanism for maintaining circuit homeostasis in response to neuronal activity (Lee et al., 2021), glial pruning is tightly regulated in a glomerulus-specific manner.

**Figure 3.**
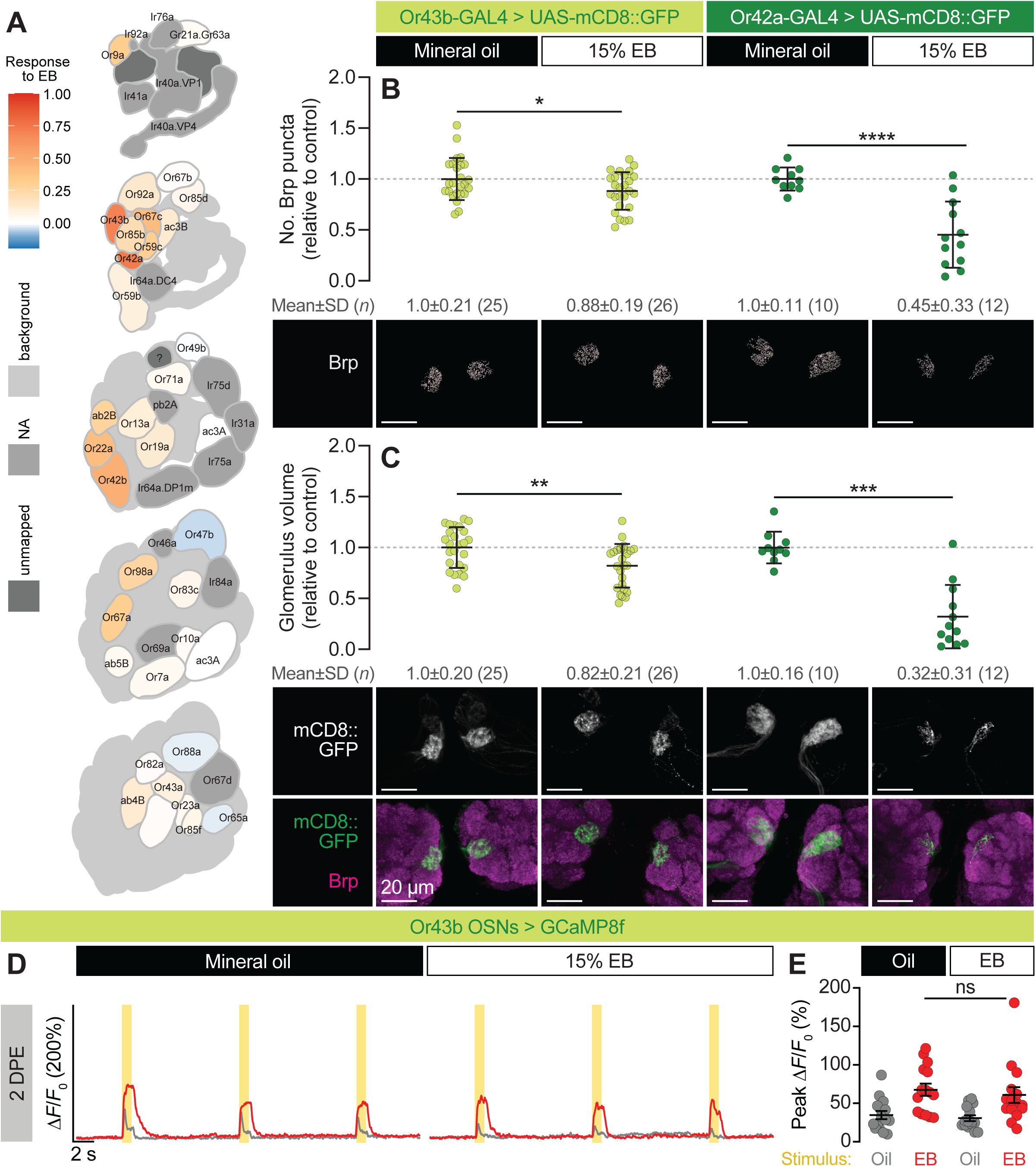
Experience-dependent pruning and inhibition are decoupled in Or43b OSNs. (**A**) DoOR 2.0 profile of EB responses across all *Drosophila* ORs. (**B**–**C**) Representative images (bottom) and quantification of presynaptic content (**B**) and volume (**C**) of VM2 (Or43b OSNs) and VM7 (Or42a OSNs) in flies exposed to mineral oil or 15% EB from 0–2 DPE. Presynapses were visualized with Brp staining. Data are mean±SD. *p<0.05, **p<0.01, ***p<0.001, ****p<0.0001, unpaired t-test. (**D**–**E**) Two-photon imaging of GCaMP8f responses in Or43b OSNs at 2 DPE following exposure to 15% EB or mineral oil from 0–2 DPE. Awake flies were stimulated with three 1 s odor puffs (yellow bars) of 5% EB (red traces and data points) or mineral oil (grey traces and data points), spaced 10 s apart. Traces (**D**) represent the mean ΔF/F_0_ values of all trials. Data points (**E**) represent the mean peak ΔF/F_0_ values for each animal. Genotypes, raw values, and detailed statistics are provided in Figure 3—source data 1.

Next, we tested the functional consequences of critical-period EB exposure on Or43b OSNs by keeping Or43b-GAL4>UAS-GCaMP8f flies in 15% EB or mineral oil from 0–2 DPE, and repeating the experimental scheme in Figure 1B. Given the inhibition observed in other EB-sensitive glomeruli, such as DM2 and DM5, following early-life EB exposure (Das et al., 2011), and our own observation of life-long Or42a OSN inhibition following glial pruning, we postulated that Or43b OSNs would be also be inhibited to some degree. Instead, we observed similar levels of EB-evoked activity in pruned Or43b OSNs relative to flies exposed to mineral oil (Figure 3D–E), suggesting that there is a nonlinear relationship between levels of pruning and OSN activity. It is possible that reduced OSN responses occur in parallel with, but not as a result of, glial pruning, or that some threshold of pruning must be reached in order for odor-evoked responses to decrease.

### Differential patterns of LN innervation may contribute to interglomerular differences in experience-dependent glial pruning

What factors drive the differences in structural remodeling and inhibition across different glomeruli during early-life EB exposure? One potential explanation is the tuning of the chemoreceptor proteins expressed by each OSN type; however, Or42a and Or43b have very similar response profiles (Grabe et al., 2016; Münch and Galizia, 2016) yet are differentially impacted by EB exposure (Figure 3). We therefore asked whether there are any prominent differences in the network connectivity of OSNs innervating VM2 and VM7, using the FlyWire full-brain connectome dataset (Dorkenwald et al., 2024; Schlegel et al., 2024; Zheng et al., 2018) to compare the upstream and downstream synaptic partners of Or24a (Figure 4A, B) and Or43b (Figure 4A, C) OSNs. Although VM7 and VM2 are innervated by relatively similar numbers of OSNs (33 and 37, respectively) and PNs (2-3 per hemisphere) (Figure 4A), VM7 OSNs far exceeded VM2 in the number of synaptic outputs and inputs as well as total number of synaptic partners, such that any given upstream or downstream partner made roughly twice as many synapses with VM7 OSNs relative to VM2 OSNs (Figure 4B–C). Strikingly, the combined volume of VM7 OSNs was ∼2.3 times that of VM2 (4,275 µm^3^ relative to 1,870 µm^3^), which was consistent with the ratio of ∼2.6 in our glomerular analysis (1,883 µm^3^ [VM7] relative to 712 µm^3^ [VM2]) (Figure 3C) and indicated that individual VM7 OSNs have greater synaptic connectivity relative to VM2 OSNs.

**Figure 4.**
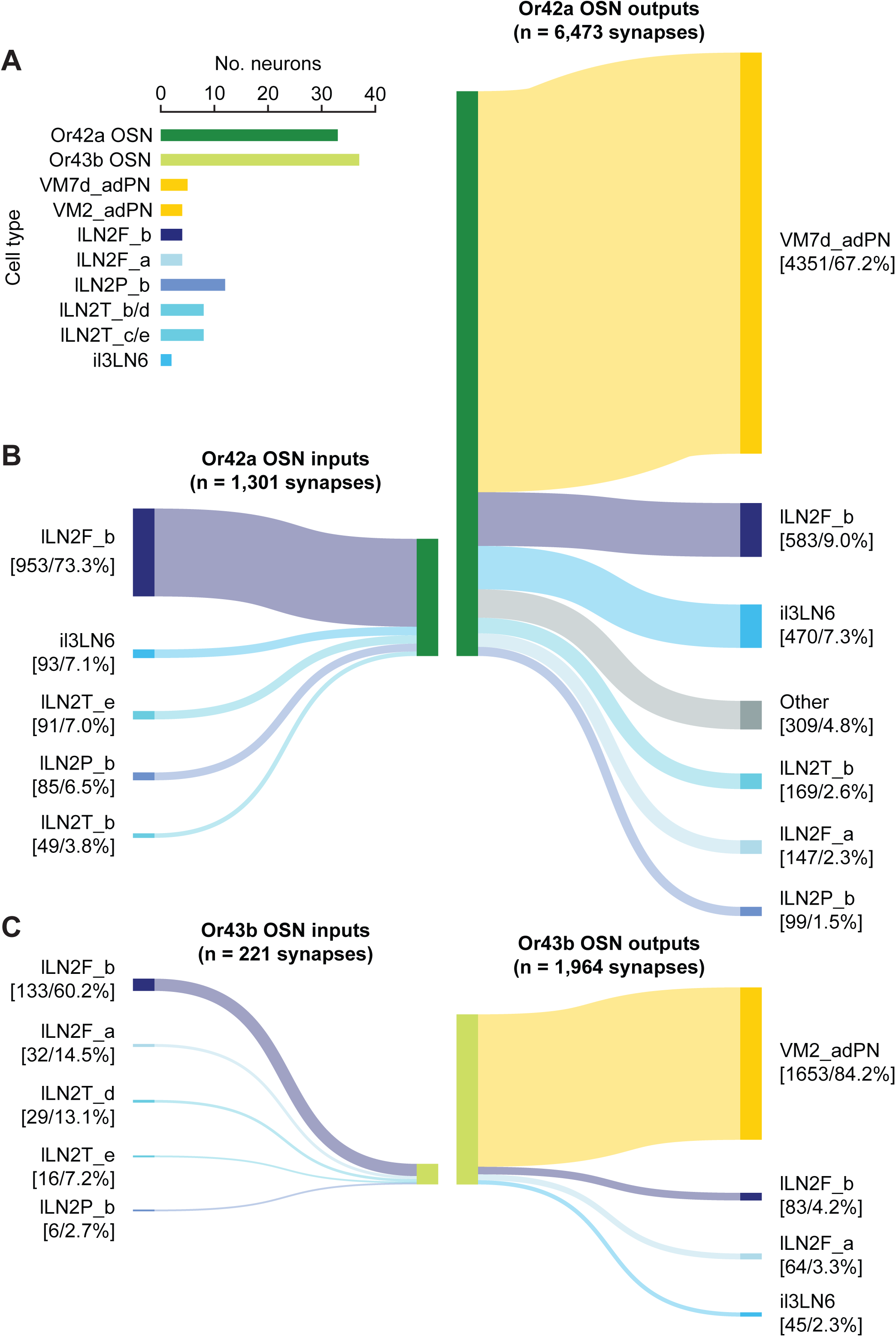
Differential patterns of LN innervation may contribute to interglomerular differences in experience-dependent glial pruning. (**A**) Numbers of neurons within the cell types included in **B** and **C**. (**B**–**C**) Sankey diagrams of the synaptic inputs and outputs of Or42a (**B**) and Or43b (**C**) OSNs. Inputs were thresholded at 99% of all synapses and outputs were thresholded at 95% of all synapses. Brackets contain the number of synapses and percentage of total synapses for that category. All connectomic analyses are derived from the FlyWire FAFB dataset. Synapse counts are provided in Figure 4—source data 1.

Both VM7 and VM2 OSNs receive the vast majority of their synaptic input from several LN subtypes that serve a variety of computational functions within the AL. For instance, both OSN types receive input from the lLN2Ps which are non-spiking, peptidergic LNs that exert intraglomerular gain control (Barth-Maron et al., 2023; Schenk and Gaudry, 2023; Sizemore et al., 2023). However, the demographics of input from LNs was not identical across Or42a and Or43b OSNs. For instance, only the Or42a OSNs received input from the bilaterally projecting il3LN6s which play a critical role in odor localization due to their asymmetric connectivity (Taisz et al., 2023). However, the most prominent difference in the connectivity of Or42a and Or43b OSNs was in their relationship with the lLN2F_b LNs. Although there are only two lLN2F_b LNs in each AL, they are a significant source of GABAergic input within the AL, collectively synapsing upon thousands of downstream targets. Due to their high degree of reciprocal connectivity within the AL, the lLN2F_bs are “rich-club” neurons (Dorkenwald et al., 2024), and have been shown to play an important role in interglomerular gain control by directly targeting OSNs (Barth-Maron et al., 2023). Strikingly, Or42a OSNs receive 953 synapses from lLN2F_b LNs, whereas Or43b OSNs receive only 133 synapses. Furthermore, while both OSN types primarily synapsed upon uniglomerular PNs, Or42a OSNs provided 583 synapses to the two lLN2F_bs, whereas Or43b OSNs only provided 83 synapses. Thus, on both the upstream and downstream side, Or42a OSNs make roughly seven-fold more inhibitory synapses with ILN2F_b inhibitory interneurons than do Or43b OSNs.

To compare the importance of GABAergic input to experience-dependent pruning of Or42a and Or43b OSNs, we performed RNAi knockdown of the key ionotropic GABA_A_ receptor Rdl (Das et al., 2011; Sudhakaran et al., 2012) in both cell types during critical-period EB or mineral oil exposure. Intriguingly, *rdl* knockdown modestly increased glial pruning of both Or42a and Or43b presynaptic terminals to a similar degree (Figure 5A–B). While this result further validates that glial pruning is proportional to EB-evoked activity, it suggests that large differences in GABAergic inhibition between Or42a and Or43b OSNs are not solely responsible for their different levels of glial pruning.

**Figure 5.**
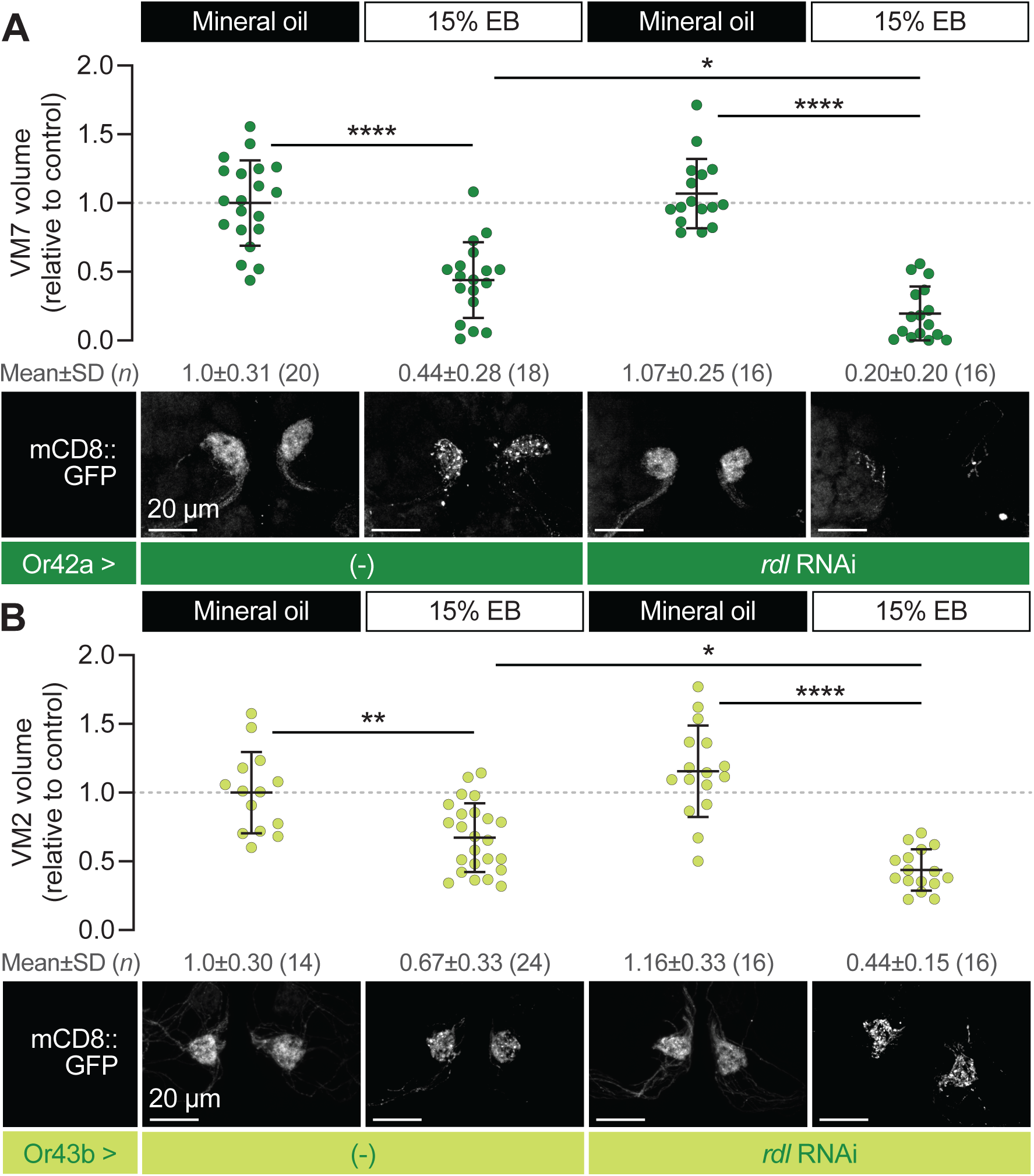
GABAergic inputs restrain glial pruning. (**A**–**B**) Representative images (bottom) and quantification of VM7 (**A**) and VM2 (**B**) volume in flies exposed to mineral oil or 15% EB from 0–2 DPE. Data are mean ± SD. *p < 0.05, **p < 0.01, ****p < 0.0001, one-way ANOVA. Genotypes, raw values, and detailed statistics are provided in Figure 5—source data 1.

LNs have been demonstrated to regulate experience-dependent plasticity in the AL, with particular insights being gained into the glomerulus-specific dynamics of GABAergic inhibitory LNs (Chodankar et al., 2020; Das et al., 2011; Mallick et al., 2024; Sachse et al., 2007). With the accessibility of detailed connectomes of the AL (Dorkenwald et al., 2024; Schlegel et al., 2024), it is possible to explore how differences in the connectivity of specific LN types to OSNs innervating each glomerulus contributes to critical period plasticity. Although there are roughly equal numbers of Or42a and Or43b OSNs and these OSN types are equally sensitive to EB, distinct patterns of connectivity could explain the differences in susceptibility to critical period pruning. The high degree of connectivity of Or42a OSNs (via the lLN2F_b LNs) could make them a “hub” of network activity during EB responses such that glia are responding to the local excitatory-inhibitory ratio, which is influenced by critical LN subtypes (Figure 4A).

In summary, we report two novel advances from our initial findings and previous studies of experience-dependent plasticity in the *Drosophila* olfactory circuit: (i) true life-long suppression of a class of OSNs following glial pruning during a critical period, and (ii) glial pruning and loss of odor sensitivity in OSNs may occur independently of each other. Importantly, these mechanisms may be viewed as part of an integrated circuit response to a sensory stressor, e.g., an overactive sensory neuron, with the end result of maintaining the odor-encoding space (Hallem and Carlson, 2006). We propose that during sensory critical periods, glial pruning and GABAergic inhibition synergize to limit network excitation and preserve odor encoding. Our work underscores the remarkable capacity of the young olfactory system to adapt to changing environmental conditions, and further positions glia as integral components of neuronal circuits.

## Methods

### *Drosophila melanogaster* stocks

Flies were reared on standard cornmeal-molasses-agar medium in a 25 °C incubator on a 12 hr/12 hr light/dark cycle, conditions which were maintained throughout odorant exposure. Unless otherwise noted, iso31 flies were used as wild-type controls in experimental crosses. Genotypes and reference information for all transgenic fly lines used in this study are provided in the Key Resources Table.

### Odorant exposure

The odorant exposure protocol was maintained from our previous study (Leier and Foden et al., 2025). For critical period exposure (0–2 days post-eclosion), 10 days after beginning crosses in standard bottles of fly food stoppered with cotton plugs, all adults (parents and any progeny that had eclosed) were anesthetized briefly with CO_2_ and discarded, and the cotton plugs were replaced with a layer of nylon mesh secured with rubber bands. A 1 mL solution of ethyl butyrate (Sigma-Aldrich #E15701) diluted volumetrically in mineral oil (Item Number (DPCI): 245-05-0562), or 1 mL of mineral oil-only control, was prepared in a 1.5 mL microcentrifuge tube, vortexed for 5–10 s, and the open tube was taped to the side of a 3.55 L airtight glass container (Glasslock #MHRB-370). The nylon-covered bottles were placed into the Glasslock containers, which were sealed and returned to the 25°C incubator. After 24 hr, all adults that eclosed during that time were anesthetized with CO_2_ and separated by genotype into standard vials of fly food secured with nylon mesh. Vials were returned to the Glasslock containers with fresh odorant solution and placed in the incubator for an additional 48 hrs, during which the odorant solution was refreshed after 24 hrs.

### Immunohistochemistry

Adult flies were anesthetized with CO_2_, decapitated, then dissected in ice-cold PBS. Once dissected, brains were fixed in ice-cold 4% (v/v) PFA in PBST (1% Triton X-100, 0.01 M phosphate, 0.0027 M KCl, and 0.137 M NaCl) for 20 minutes. Brains were immediately washed three times with PBST, incubated for 1 hr in blocking buffer (5% normal goat serum, 0.3% Triton X-100 in PBS), then incubated for 24 hrs at 4°C with primary antibodies diluted in blocking buffer. Brains were given three 10 min washes with PBST, then incubated for 24 hrs at 4°C with secondary antibodies diluted in blocking buffer as before. The next day, brains were given three 10 min washes with PBST and kept stationary for 30 min in SlowFade Gold mountant (Thermo Fisher #S36936). Mounting was performed essentially as described (Kelly et al., 2017). Unless otherwise noted, all steps were performed with rocking at room temp. A complete list of antibodies and dilutions is provided in the Key Resources Table.

### Confocal microscopy

Images were acquired with a Zeiss LSM 800 confocal laser scanning microscope using a 100 x/1.4 NA Plan-Apochromat objective. Laser intensity and gain were optimized for each z-stack.

### Image analysis

Quantification of glomerulus volume and presynaptic content was done using Imaris 9.7.1 (Bitplane). First, the Surfaces function was used on the GFP channel to model the OSN membrane. The following settings were used: baseline subtraction (threshold of 1000) followed by background subtraction (filter width of 10 µm); smoothing enabled (surfaces detail of 0.1 µm); thresholding based on local contrast (largest sphere diameter set to 10 µm), with automatic thresholding enabled; and filter based on number of voxels (cutoff of 500 voxels). The resulting Surfaces were manually processed by a blinded experimenter who used the scissors tool to remove OSN axons prior to the terminal arbor and used the final Surfaces object to mask the Brp channel. The Spots function was then used on the masked channel to quantify the number of Brp puncta within the glomerulus, using the following settings: deconvolution (robust algorithm, standard parameters) followed by background subtraction (filter width of 10 µm); algorithm set to different Spots sizes (region growing); background subtraction enabled; estimated diameter of 0.4 µm; automated quality filter; Spots regions from absolute intensity, with automatic region thresholding enabled.

### Fly preparation for *in vivo* Ca^2+^ imaging and odor delivery

All *in vivo* Ca^2+^ imaging experiments were performed immediately following odorant exposure using a Bruker Ultima two-photon microscope system (Bruker Corporation, Billerica, MA) equipped with a Ti:Sapphire laser (Coherent Inc., Santa Clara, CA), tuned to 920 nm for imaging and controlled by Prairie View software (Bruker). Fluorescence signals were detected using a gallium arsenide phosphide (GaAsP) photomultiplier tube. Imaging was performed at a frame rate of 29.3 Hz. Both male and female flies were used.

Flies were anesthetized on ice and transferred to a custom recording chamber consisting of a square aluminum foil sheet (10 mm × 12 mm) affixed to a plastic Petri dish. A central imaging window (∼1 mm × 1 mm) was cut into the foil to expose the head. Flies were positioned in the window and secured using a UV-cured plastic welder (BONDIC SK8024, NY). The head was immobilized such that the antennae remained dry during dissection. A small incision was made in the cuticle using 26-gauge needles (BD PrecisionGlide, 305110-26G), and overlying tissue was removed to expose the dorsal surface of the brain for imaging.

The Or42a-GAL4 and Or43b-GAL4 driver lines (Key Resources Table) were used to visualize olfactory sensory neurons (OSNs) projecting to their respective glomeruli. For Ca²⁺ imaging experiments, GCaMP8f was expressed in OSNs to measure odor-evoked responses to Ethyl Butyrate. For voltage imaging, ASAP5 was expressed in OR42a OSNs. Imaging was performed in the antennal lobe, with a particular focus on glomerular responses to odor stimulation. The imaging chamber was filled with approximately 3 mL of saline.

For odor stimulation, ethyl butyrate was diluted 5:100 in mineral oil and delivered using a modified version of a previously described method. In brief, 50 µL of diluted odorant (in mineral oil) was pipetted onto a small piece of Whatman filter paper, which was inserted into the wide end of a 5¾-inch glass Pasteur pipette (Chemglass Life Sciences, Cat. No. CG-1700-14, Fisher Scientific). The tapered tip of the pipette was placed into a glass stimulation tube that received a continuous humidified air stream. The Pasteur pipette was positioned such that a lateral opening in the stimulation tube allowed for the integration of the odorized airstream into the main flow path. An air source was connected to the back of the pipette, behind the filter paper, enabling airflow to push volatile odorants through the pipette and out the tip into the stimulation tube. Odorized airflow was regulated via a carbon-filtered compressed air system split into a constant background stream (2.5 L/min; Dwyer VFA-25-BV flowmeter) and an odor stream (0.8 L/min; Dwyer VFA-23-BV flowmeter). A Parker solenoid valve (001-0028-900, Hollis, NH) toggled between the odor pipette and constant air flow. Odor stimuli were delivered for 1 second, three times per trial using Prairie View software.

Raw imaging data were imported into FIJI for ROI selection. Fluorescence signals were extracted and processed using a custom Matlab script. Signal changes were normalized to the baseline fluorescence (F), calculated as the average over the 3 seconds preceding the first odor stimulus. ΔF/F was calculated as the percent change from baseline, and traces and summary data were visualized accordingly. Processed data were analyzed statistically using GraphPad Prism.

### Connectomic analysis

FlyWire (Dorkenwald et al., 2024; Schlegel et al., 2024) analysis was performed with the fafbseg (Dorkenwald et al., 2024; Schlegel et al., 2023; Zheng et al., 2018) python package. VM7 and VM2 OSN presynaptic and postsynaptic connectivity was extracted using get_connectivity and NeuronCriteria functions in the flywire module of fafbseg. The presynaptic and postsynaptic root IDs collected were then matched with their respective cell types from get_heirarchical_annotations in the flywire module of fafbseg. The data were filtered to exclude connections with less than five synapses. We sorted the inputs to VM7 OSNs and VM2 OSNs and thresholded the connectivity at 99% of total synaptic partners. We thresholded VM7 OSN and VM2 OSN outputs at 95% of total partners because they have at least four times as many postsynapses compared to presynapses. “Other” consisted of all cells that were not annotated as lLN2F_b/a, lLN2P_b, lLN2T_b/c/d/e, il2LN6 or PN.

### Statistics

Statistical analyses were performed with Prism 10.2.1 (GraphPad). All data is from at least two independent experiments. Sample size was not predetermined. For microscopy experiments, each data point represents one glomerulus; for calcium imaging experiments, each data point represents recordings from one animal. Shapiro-Wilks normality testing showed that our data was not normally distributed, so nonparametric tests were used: the Mann-Whitney test for comparisons of two samples, and the Kruskal-Wallis test with Dunn’s test for multiple comparisons for three or more samples. All tests were two-tailed with an alpha of 0.05, and no outliers were excluded. p-values are represented as follows: ns, not significant (p≥0.05); *p<0.05; **p<0.01; ***p<0.001, ****p<0.0001.

## Acknowledgements

We thank the Princeton FlyWire team and members of the Murthy and Seung labs, as well as members of the Allen Institute for Brain Science, for development and maintenance of FlyWire (supported by BRAIN Initiative grants MH117815 and NS126935 to Murthy and Seung). We also acknowledge members of the Princeton FlyWire team and the FlyWire Consortium for neuron proofreading and annotation. We thank Jaeda Coutinho-Budd, Dan Jindal, and members of the Broihier and Dacks labs for thoughtful feedback and stimulating conversations. We gratefully acknowledge stocks obtained from the Janelia Research Campus and the Bloomington Stock Center (NIH P400D018537). This work was supported by NIH grants F30AG097357 (HCL), R01DC016293 (AMD, JJ, AJW), T32GM152319 (HCL, AJF), and R01NS120689 (HCL, AJF, PVLC, HTB).

